# SIX1 cooperates with RUNX1 and SMAD4 in cell fate commitment of Müllerian duct epithelium

**DOI:** 10.1101/427351

**Authors:** Jumpei Terakawa, Vanida A. Serna, Devi Nair, Shigeru Sato, Kiyoshi Kawakami, Sally Radovick, Pascal Maire, Takeshi Kurita

## Abstract

During female mammal reproductive tract development, epithelial cells of the lower Müllerian duct are committed to become stratified squamous epithelium of vagina and ectocervix, when the expression of ΔNp63 transcription factor is induced by mesenchymal cells. The absence of ΔNp63 expression leads to adenosis, the putative precursor of vaginal adenocarcinoma. Our previous studies with genetically engineered mouse models have established that fibroblast growth factor (FGF)/mitogen-activated protein kinase (MAPK), bone morphogenetic protein (BMP)/SMAD, and activin A/runt related transcription factor 1 (RUNX1) signaling pathways are independently required for ΔNp63 expression in Müllerian duct epithelium (MDE). Here we report that sine oculis homeobox homolog 1 (SIX1) plays a critical role in the activation of ΔNp63 locus in MDE as a downstream transcription factor of mesenchymal signals. In mouse developing reproductive tract, SIX1 expression was restricted to MDE of the future cervix and vagina. SIX1 expression was totally absent in SMAD4 null MDE and was reduced in RUNX1 null and FGFR2 null MDE, indicating that SIX1 is under the control of vaginal mesenchymal factors, BMP4, activin A and FGF7/10. Furthermore, *Six1, Runx1* and *Smad4* gene-dose-dependently activated ΔNp63 expression in MDE within vaginal fornix. Using a mouse model of diethylstilbestrol (DES)-associated vaginal adenosis, we found DES action through epithelial estrogen receptor α (ESR1) down-regulates SIX1 and RUNX1 in MDE within the vaginal fornix. This study establishes that the vaginal/ectocervical cell fate of MDE is regulated by a collaboration of multiple transcription factors including SMAD4, SIX1 and RUNX1, and the down-regulation of these key transcription factors leads to vaginal adenosis.

**Author Summary:** In embryogenesis, differentiation fate of cells is specified through constant communication between neighboring cells. In this study, we investigated the molecular mechanism of epithelial cell fate commitment in the lower female reproductive organs utilizing mouse genetic models. The cell fate of epithelial cells in the uterus, cervix and vagina is directed by signaling from mesenchymal cells. We demonstrated that within the epithelial cells of the developing vagina, signals from mesenchymal cells are integrated into activities of transcription factors including SMAD4, RUNX1 and SIX1, which dose-dependently co-operate in the determination of vaginal epithelial cell fate. Disruption of these processes alters the cell fate from vaginal to uterine epithelium, resulting in a condition called vaginal adenosis, a putative precursor of vaginal adenocarcinoma. Women exposed to diethylstilbestrol (DES) in the womb have about 40 times the risk of developing vaginal adenocarcinoma. We determined that developmental exposure to DES induces vaginal adenosis by repressing SIX1 and RUNX1 through ESR1 in the epithelial cells. This discovery enhances the understanding of how early-life events, such as exposure to endocrine disruptors, causes vaginal adenosis, and thus may contribute to the prevention and therapeutic treatment of idiopathic vaginal adenocarcinoma.

## INTRODUCTION

In mammals, the majority of female reproductive tract (FRT) develops from the Müllerian ducts (MDs) [1-3]. During embryogenesis, the MDs undergo a dynamic transformation from simple tubes consisting of homogeneous epithelium and mesenchyme into distinct organs, namely the oviduct, uterus, cervix and vagina [2, 4]. Classic tissue recombination studies have established that organ-specific mesenchyme induces the differentiation of MD epithelium (MDE) into epithelia with unique morphology and functions [5-7]. In the lower MD, epithelial cells are committed to become stratified squamous epithelium of ectocervix and vagina (together referred to as “vagina” hereafter), as the expression of ΔNp63 transcription factor is induced by vaginal mesenchyme [8-10]. In MDE of the developing vagina, the expression of ΔNp63 is activated by mesenchymal paracrine factors: bone morphogenetic protein (BMP) 4, activin A (ActA) and fibroblast growth factor (FGF) 7 or 10 [11, 12]. SMAD4 is essential for the activation of ΔNp63 in MDE, and this transcription factor binds on the 5’ sequence adjacent to the transcription start site (TSS) of ΔNp63 in future vaginal epithelium (VgE) but not in future uterine epithelium (UtE) [12]. This SMAD-dependent activation of the ΔNp63 locus requires runt-related transcription factor 1 (RUNX1), a co-transcription factor of SMADs. In MDE, the expression of RUNX1 is activated by ActA through a SMAD-independent mechanism [11]. In addition, activation of the mitogen-activated protein kinase (MAPK) pathway by FGF7/10-FGF receptor 2 IIIb (FGFR2IIIb) is essential for the activation of ΔNp63 locus in MDE [11]. BMP4-SMADs, ActA-RUNX1 and FGF7/10-MAPK pathways are independently required for the vaginal cell fate commitment of MDE, as inactivation of *Smad4, Runx1* or *Fgfr2* in MDE results in uterine epithelial differentiation of MDE within the vagina, which is a congenital epithelial lesion called vaginal adenosis [11, 12]. Nevertheless, once the ΔNp63 locus is activated in MDE, the transcriptional activity of the ΔNp63 locus is cell-autonomously maintained by ΔNp63 protein itself [12]. Hence, the identity of VgE is maintained independent of mesenchymal factors [7, 8].

In this study, we investigated the role of sine oculis homeobox homolog 1 (SIX1) in the cell fate commitment of VgE. In mammals, *SIX1* and other five *SIX* genes (*SIX2–6*) synergistically regulate the developmental process in multiple organs, including inner ear, salivary gland, kidney, lung, and trachea [13, 14]. In mouse FRTs, *Six1* is enriched in the vagina compared to the uterus [12, 15]. However, its biological function in FRT remains unclear. Our current mouse genetic study reveals that SIX1 co-operates with RUNX1 and SMAD4 in the activation of the ΔNp63 locus in MDE as a downstream transcription factor of BMP4, ActA and FGF7/10 in MDE. The etiology of vaginal adenosis, the putative precursor to vaginal adenocarcinoma (VAC) is commonly associated with intrauterine exposure to estrogenic compounds, including diethylstilbestrol (DES) [16]. Our previous studies established that DES induces vaginal adenosis through inhibition of ΔNp63 expression in MDE. Our current study provides evidence that DES blocks the activation of ΔNp63 locus in future VgE by repressing SIX1 and RUNX1 through epithelial estrogen receptor *α* (ESR1). Such discoveries from our models may contribute to the prevention and therapeutic treatment of VACs, the etiology of which is currently unknown.

## RESULTS

### Expression patterns of SIX1 in neonatal FRTs

ΔNp63α is the dominant isoform of the transcription factor encoded by *Trp63/TP63* in mouse/human VgE [10, 12]. To identify molecules that control epithelial cell fate in the lower FRT, we conducted microarray analysis of neonatal vagina and uterus from MDE-specific conditional KO (cKO) and conditional heterozygous (cHet, control) mice of *Trp63* [12]. In the analysis, *Six1* was more enriched in vaginae than uteri (1.02 Log2 fold-change, p = 0.0013) at postnatal day 2 (PD2), when induction of ΔNp63 expression is in progress in the vagina (Fig 1A). The expression level of *Six1* transcripts was not significantly different between *Trp63* cKO and cHET mice (Log2 cHET/cKO = −0.176, p = 0.23) (GSE44697) [12], indicating that SIX1 is not the target of TRP63. Immunoblotting confirmed the results of microarray: SIX1 protein was detected in vaginae but not in uteri and ovaries from PD2 C57BL/6J mice (Fig 1B).

**Fig 1.**
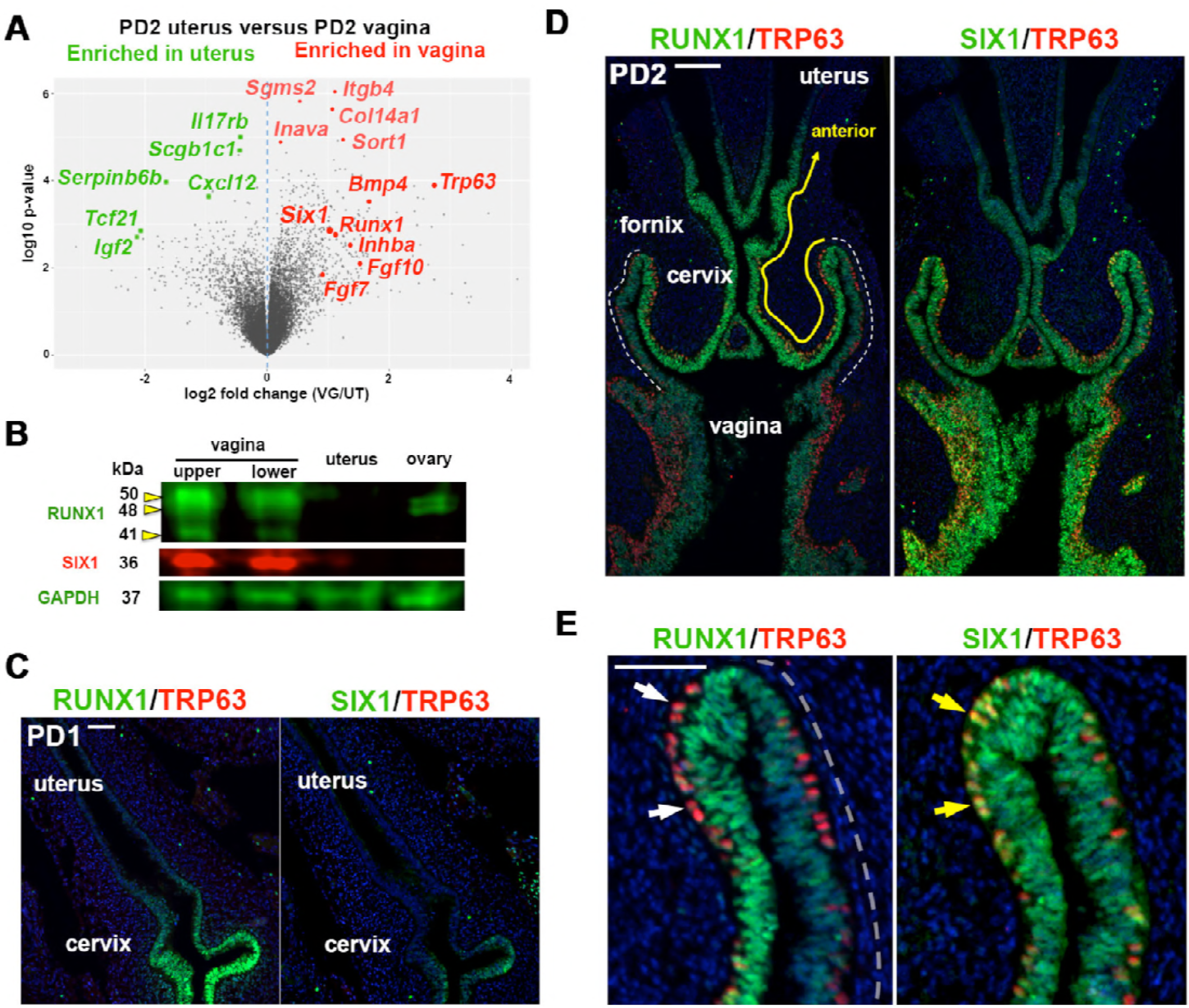
Expression patterns of SIX1 in developing female reproductive tract. (A) Volcano plot displaying differential expressed genes in mouse PD2 uterus and vagina. Genes significantly enriched in vagina and uterus in microarray analysis [12] are marked in red and green, respectively. (B) Immunoblot analysis of SIX1 and RUNX1 expression in PD2 mouse FRT. The vagina was divided into upper and lower half. (C-E) Immunofluorescence assay for RUNX1, SIX1 (green) and TRP63 (red) in the lower FRT of PD1 (C) and PD2 (D, E) mice. Outer–wall of fornix is marker with dotted line (D). In the vaginal fornix (E), RUNX1 is down-regulated in MDE upon expression of TRP63 (white arrows), whereas TRP63 and SIX1 are co-expressed (yellow arrows). Bar = 100 μm (C and D), = 50 μm (E).

Similarly to the expression of ΔNp63 in developing vagina, SIX1 expression progressed from posterior to anterior. At birth, SIX1 was expressed in the MDE of the lower vagina but not in the upper vagina and cervix, where RUNX1 already highlighted the future VgE (Fig 1C). By PD2, SIX1 expression extended to the cervix (Fig 1D), thus SIX1 and RUNX1 were co-expressed in the future VgE. There were substantial differences in the expression patterns of SIX1 and RUNX1 in neonatal FRTs. RUNX1 was concentrated in the MDE in the cervical canal and the upper-portion of vagina, and the expression was reduced in the posterior portion from the outer-wall of the fornix (Fig 1D, outer-wall of fornix is marked with white dotted-line), whereas SIX1 was expressed at a similar level in both inner and outer walls of the fornix (Fig 1D). In addition, RUNX1 in MDE was down-regulated upon expression of ΔNp63 (Fig 1E, white arrow) [12], whereas SIX1 expression persisted in ΔNp63 positive cells (Fig 1E, yellow arrow).

### SIX1 is a downstream transcription factor of SMAD4

SIX1 was expressed in the fornices of ΔNp63 cKO and cHET mice [12] at PD14, confirming that expression of SIX1 is independent of ΔNp63 (Fig 2A). In contrast, expression of SIX was SMAD4 dependent: *Smad4* cKO mice [12] completely lacked the expression of SIX1 in the entire MDE as assessed at PD2 (n = 5) (Fig 2B).

**Fig 2.**
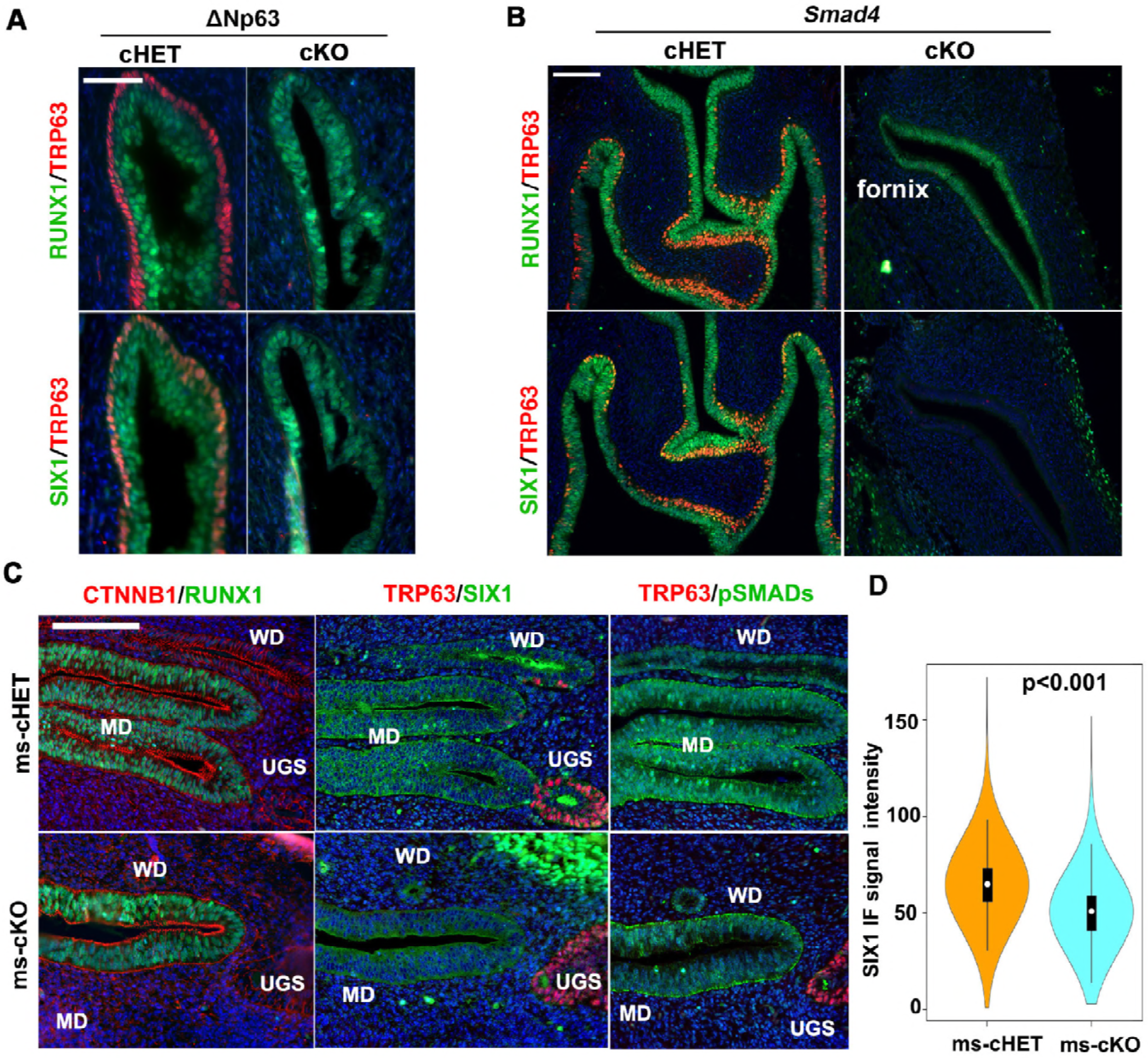
SIX1 is a down-stream transcription factor of BMP4-SMAD pathway. In all figures, outer-wall of fornix is shown on the right side. (A) SIX1 expression is maintained in the vaginal fornix of ΔNp63 cKO mice (PD14) (n ≥ 4). Bar = 50 μm. (B) SIX1 expression in MDE is SMAD4 dependent. At PD2, SIX1 is totally absent in the MDE of *Smad4* cKO mice, which normally express RUNX1 in MDE. Bar = 100 μm. (C) Deletion of *Bmp4* in mesenchymal cells reduces SIX1 in MDE. (D) Violin plot of SIX1 immunostaining signals in the lower MDE of *Bmp4* ms-cHET and ms-cKO mice (n = 3 each). The signal distributions of two groups are significantly different (p<0.01). Bar = 100 μm.

The absence of SIX1 in *Smad4* cKO mice suggested that SIX1 is the downstream transcription factor of BMP4-SMAD pathway. Therefore, *Bmp4* was knocked out in vaginal mesenchyme utilizing *Twist2^Cre^* [17], and the effect on the expression of SIX1 in the lower Müllerian duct was assessed. Mesenchyme-specific *Bmp4* conditional KO (ms-cKO) was embryonic lethal. Hence, we collected pelvic organs from *Bmp4* conditional ms-cKO and ms-cHET mice at embryonic day 15.5, when most *Bmp4* ms-cKO embryos exhibited normal growth. At embryonic day 15.5 (Fig 2C), RUNX1 expression highlighted the anterior portion of MDE in both *Bmp4* ms-cKO and ms-cHET mice. On the other hand, SIX1 expression in MDE was low and mostly cytoplasmic at this age. Nevertheless, the SIX1 signal in MDE was higher in *Bmp4* ms-cHET mice compared to *Bmp4* ms-cKO mice (n=3 each, Fig 2D), suggesting that BMP4 is a factor regulating SIX1 expression in FRT. However, the effect of loss of BMP4 on SIX1 expression was not comparable to the loss of SMAD4. This might be due to incomplete deletion of *Bmp4* or compensation by other BMP family members, as phosphorylation of SMAD1/5/9 was still present in the MDE of *Bmp4* cKO mice (Fig 2C).

### ActA-RUNX1 and FGF7/10-MAPK pathways positively regulate SIX1 in MDE

Although SIX was expressed throughout the VgE in *Runx1* cKO mice [12] (Fig 3A), the expression of SIX1 in the fornix was significantly reduced compared to *Runx1* cHET mice (Fig 3B and 3C). Thus, the expression level of SIX1 in MDE is positively regulated by ActA-RUNX1 signaling activity. Similarly, SIX1 expression was slightly reduced in the fornix of *Fgfr2* cKO mice [11] (Fig 3D). However, SIX1 expression in the fornix was uniformly up-regulated when the vaginal defect of *Fgfr2* cKO MDE was corrected with the expression of a constitutively active MAP2K1 (MAP2K1^DD^) [11] (Fig 3D and 3E), suggesting that MAPK activity modulates the expression level of SIX1 protein in the vaginal fornix. Accordingly, we tested the effect of BMP4, ActA and FGF10 on SIX1 expression in uterine organ culture assay. ActA and FGF10 had minimal to no effect on SIX1 expression in the epithelium of uterine explants (not shown). BMP4 slightly increased SIX1 in UtE, but the nuclear expression was mostly absent (Fig 3F). Even when all 3 factors were combined, nuclear SIX1 expression was detected only in portions of UtE showing ΔNp63 expression, suggesting that SIX1 promotes ΔNp63 expression in MDE. In the uterine organ culture, growth factors in the medium must diffuse through the mesenchymal layers to act on UtE. Diffusion of FGF10 within connective tissues is limited because of its high affinity to heparan sulfate [18]. Accordingly, we replaced FGF10 with the expression of MAP2K1^DD^, which itself did not induce expression of ΔNp63, RUNX1 and SIX1 [11]. ActA and BMP4 efficiently induced SIX1 as well as ΔNp63 in *Map2k1^DD^* transgenic UtE (Fig 3F and 3G) indicating that SIX1 is the downstream transcription factor of BMP4-SMAD, ActA-RUNX1 and FGF7/10-MAPK in MDE.

**Fig 3.**
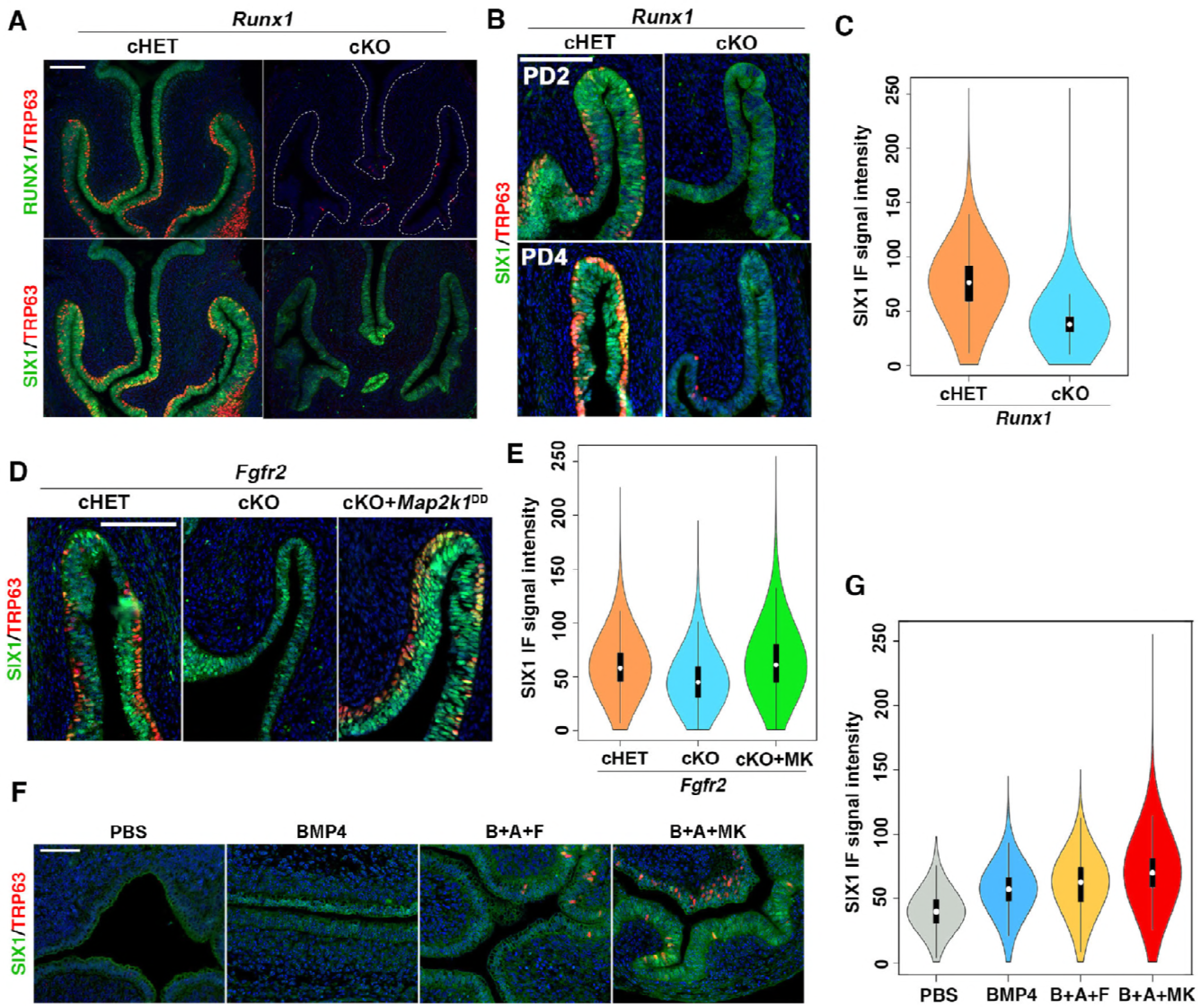
RUNX1 and FGFR2 modulate expression levels of SIX1 in MDE. (A) Expression of RUNX1 and SIX1 in the lower FTR of PD2 *Runx1* cHET and cKO mice. RUNX1 null vaginal/cervical epithelium is outlined by doted lines. Nuclear expression of SIX1 expression is reduced in the fornices of *Runx1* cKO mice. (B) SIX1 expression patterns in the vaginal fornices of *Runx1* cHET and cKO mice at PD2 and PD4. In the fornix of *Runx1* cKO mice, nuclear expression of SIX1 increases from PD2 to PD4, but the overall expression level of SIX1 in MDE remains low and uneven. (C) Violin plot of SIX1 IF signal distribution in the fornix of PD2 *Runx1* cHET and cKO mice (n ≥ 4 per group). The signal distributions of two groups are significantly different (p<0.01). (D) Expression of SIX1 in the vaginal fornix of *Fgfr2* mutant mice. SIX1 is reduced in the fornix of *Fgfr2* cKO mice, but the SIX1 expression level is restored by expression of MAP2K1^DD^. (E) Violin plot of SIX1 IF signal distribution in the fornix of PD2 *Fgfr2* cHET, *Fgfr2* cKO and *Fgfr2* cKO with MAP2K1^DD^ (cKO+MK) mice (n = 4 per group). The signal distributions are significantly different among 3 groups (p<0.01). (F) Regulation of SIX1 in cultured uterine explants. 20 ng/ml BMP4 has a weak effect on the expression of SIX1 in UtE. The combination of BMP4 (B), ActA (A) and FGF10 (F) (20 ng/ml each) induced nuclear expression of SIX1 and ΔNp63 in restricted regions of UtE. Replacement of FGF10 with *Map2k1^DD^* transgene (MK) efficiently induced SIX1 and ΔNp63 in UtE. (G) Violin plot of SIX1 IF signal distribution in the UtE of cultured uterine explants (n ≥ 4 per group). The signal distributions are significantly different among groups (p<0.01). Bars = 100 μm.

### *Six1* and *Runx1* dose-dependently promote ΔNp63 expression in MDE

Since Six1 null mice die before vaginal epithelial differentiation occurs [19], the role of SIX1 in VgE differentiation was assessed by genetically inactivating *Six1* in MDE by *Wnt7a-Cre* [20]. *Six1* cKO mice were born with the expected Mendelian ratio demonstrating no gross abnormality. However, loss of SIX1 in MDE affected the formation of ΔNp63-positive basal epithelial layer in the vaginal fornix, and a substantial area of epithelium was negative for ΔNp63-positive cells at PD4 (Fig 4A). Thus, SIX1 is one of key transcription factors that mediate the paracrine mesenchymal signaling in the vaginal cell fate commitment of MDE. Nevertheless, the defect of *Six1* cKO vagina was relatively minor, and a continuous ΔNp63 positive layer formed in the fornix by PD14, as the lateral growth of ΔNp63 positive cells filled the gaps (not shown). The distinctive vaginal phenotypes of *Six1* cKO mice from *Smad4, Runx1* and *Fgfr2* cKO mice indicate that SIX1 is only one of many downstream factors mediating signaling from mesenchymal cells in MDE. While *Smad4, Runx1* and *Fgfr2* cKO mice lost ΔNp63 expression in MDE within the entire (*Smad4*, and *Fgfr2* cKO) or upper (*Runx1* cKO) vagina, the vaginal defect of *Six1* cKO mice was restricted to the epithelium on the outer-wall of vaginal fornix, where the expression of RUNX1 is reduced (Fig 1D). Meanwhile, RUNX1 expression in the vaginal fornix was not affected in *Six1* cKO mice (Fig 4A). Hence, we generated the compound conditional mutant mice of *Six1* and *Runx1* to assess if SIX1 and RUNX1 collaborate in the ΔNp63 expression of MDE in the outer-wall of vaginal fornix. Monoallelic loss of *Runx1* in MDE exaggerated the effect of *Six1* alleleic loss on ΔNp63 expression: While monoallelic loss (cHET) of *Six1* or *Runx1* alone had no evident effect on the formation of ΔNp63-positive basal layer, the *Six1;Runx1* double cHET mice had a significantly reduced number of ΔNp63-positive basal cells in the outer-wall of fornix (Fig 4B and 4C). The ΔNp63-negative epithelial area expanded further to the inner wall of fornix when biallelic loss (cKO) of *Six1* was combined with monoallelic loss of *Runx1* (Fig 4B and 4C). The differentiation of MDE itself was not retarded in the mutant mice, as ΔNp63 negative MDE expressed progesterone receptor (PGR), indicating uterine cell fate commitment [21] (Fig 4D).

**Fig 4.**
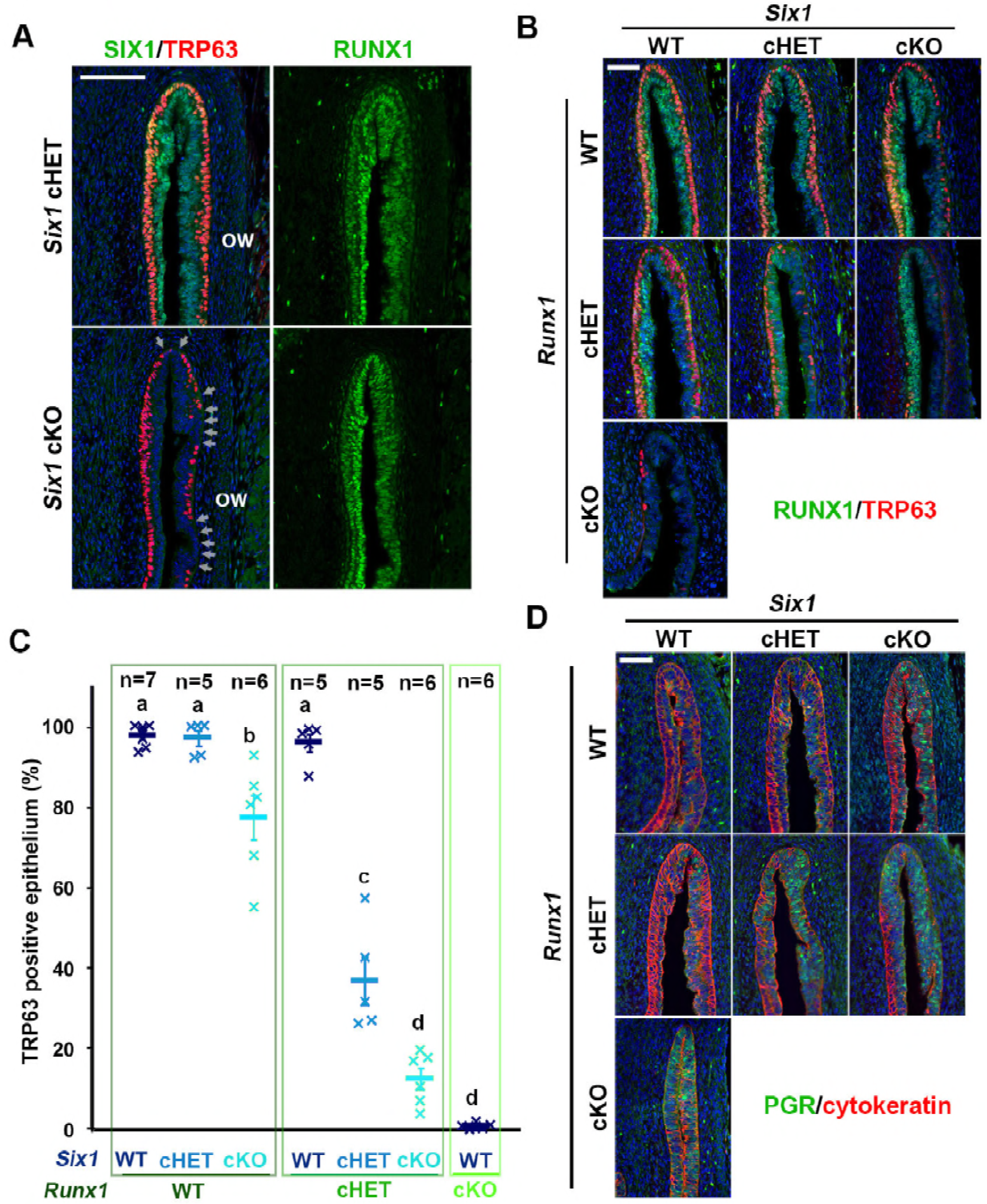
SIX1 and RUNX1 collaborate in the activation of ΔNp63 locus in MDE. (A) Six1 cKO mice showed minor defects in ΔNp63 expression in the outer-wall (ow) of vaginal fornix. The ΔNp63 negative epithelial regions are indicated by arrows. (B-D) Gene-does effect of *Six1* and *Runx1* on vaginal cell fate commitment of MDE in the vaginal fornix. The outer fornix wall is on the right side. (B) Expression of ΔNp63 (red) and RUNX1 (green). (C) Proportion of MDE lined with ΔNp63-psotive basal layer on the outer-wall of vaginal fornix. (D) Expression of uterine epithelial marker (PGR, green). The epithelium is highlighted with cytokeratin (red). Bars = 100 μm.

### Gene-dose-dependent function of *Six1, Runx1* and *Smad4* in activation of ΔNp63 locus in MDE

The distinctive vaginal phenotypes of *Six1* cKO and *Smad4* cKO mice indicate that SMAD4 works independent of SIX1 in vaginal cell fate commitment of MDE. Accordingly, we assessed if the efficacy of SIX1 and RUNX1 in the activation ΔNp63 expression in MDE is affected by monoallelic loss of *Smad4* gene, which alone does not block the formation of ΔNp63-positive basal layer in VgE [12]. *Six1;Smad4* double cHET mice expressed ΔNp63 throughout the vagina at PD4. However, the density of basal cells on the outer-wall of the fornix was reduced (Fig 5). The synergy between *Six1* and *Smad4* alleles became more prominent when an additional *Six1* allele was inactivated (Fig 5). Similarly, monoallelic loss of *Smad4* and *Runx1* synergistically affected the density of ΔNp63 in the fornix. Accordingly, *Six1;Smad4;Runx1* triple cHET mice demonstrated gaps in the ΔNp63-positive basal layer throughout the vaginal fornix (Fig 5). The effect of monoallelic *Smad4* loss on the density of TRP63 positive cells was statistically significant in mice with certain genotypes (Table 1). For instance, TRP63-positive cell density in the outer and inner fornix walls of *Six1* cHET mice was not significantly different from that in WT mice. However, the TRP63-positive cell density in *Six1* cHET mice became significantly lower with the monoallelic loss of *Smad4* (*Six1* cHET; *Smad4* cHET) compared to WT mice (Fig 5 B and C).

**Fig 5.**
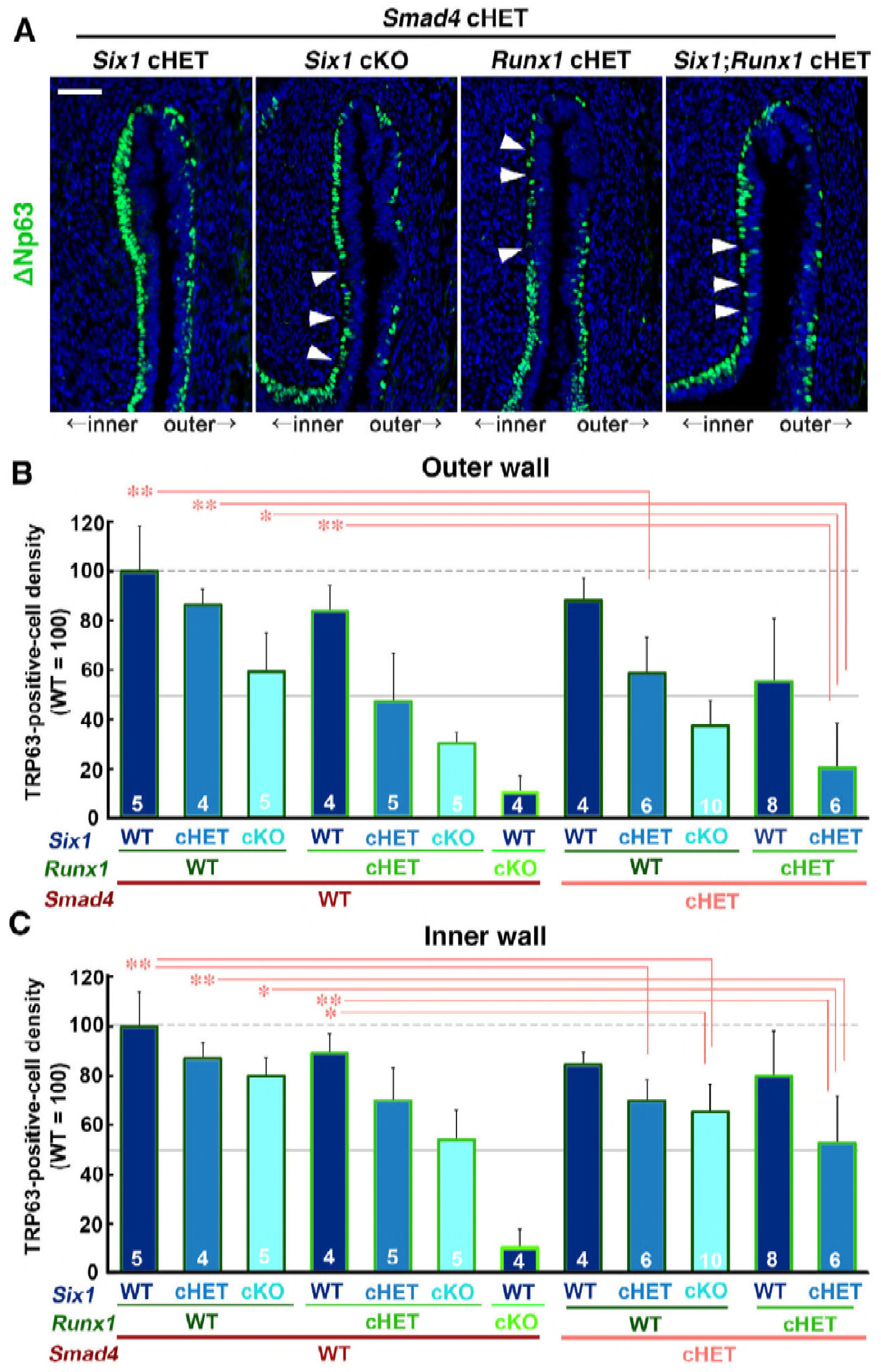
Dose-dependent function of *Six1, Runx1* and *Smad4* in the activation of ΔNp63 locus. (A) Monoallelic loss of *Smad4* exaggerates effects of *Six1* and *Runx1* null alleles on ΔNp63 expression (green) in MDE. The outer fornix wall is shown on the right side. Breaks in the ΔNp63-positive basal layer in the inner fornix wall are marked by arrowheads. Bar = 50 μm. (B and C) Basal cell density (TRP63-positive nuclear area per epithelial basement membrane length) in the outer and inner fornix walls of *Six1, Runx1* and *Smad4* compound mutant mice. The sample number in each group is marker on the bars. The result is demonstrated by average means ± SD. The comparisons that become significantly different by monoallelic *Smad4* loss are marked with lines, and the groups with significantly higher value are marked with asterisks. * p< 0.05, ** p< 0.01.

**Table 1.** One-way ANOVA followed by Tukey’s multiple comparison test of TRP63-posittive cell density in the outer and inner vaginal fornix walls of *Six1, Runx1, Smad4* compound mutant mice.

| Outer wall |  | Six1 | Smad4 WT |  |  |  |  |  |  | Smad4 HET |  |  |  |  |
| --- | --- | --- | --- | --- | --- | --- | --- | --- | --- | --- | --- | --- | --- | --- |
|  |  |  | Runx1 WT |  |  | Runx1 HET |  |  | Runx1 KO | Runx1 WT |  |  | Runx1 HET |  |
|  |  |  | WT | HET | KO | WT | HET | KO | WT | WT | HET | KO | WT | HET |
| Smad4 WT | Runx1 WT | WT |  | NS | p<0.01 | NS | p<0.01 | p<0.01 | p<0.01 | NS | p<0.01 | p<0.01 | p<0.01 | p<0.01 |
|  |  | HET |  |  | NS | NS | p<0.05 | p<0.01 | p<0.01 | NS | NS | p<0.01 | NS | p<0.01 |
|  |  | KO |  |  |  | NS | NS | NS | p<0.01 | NS | NS | NS | NS | p<0.01 |
|  | Runx1 HET | WT |  |  |  |  | p<0.05 | p<0.01 | p<0.01 | NS | NS | p<0.01 | NS | p<0.01 |
|  |  | HET |  |  |  |  |  | NS | p<0.05 | p<0.01 | NS | NS | NS | NS |
|  |  | KO |  |  |  |  |  |  | NS | p<0.01 | NS | NS | NS | NS |
|  | Runx1 KO | WT |  |  |  |  |  |  |  | p<0.01 | p<0.01 | NS | p<0.01 | NS |
| Smad4 HET | Runx1 WT | WT |  |  |  |  |  |  |  |  | NS | p<0.01 | p<0.05 | p<0.01 |
|  |  | HET |  |  |  |  |  |  |  |  |  | NS | NS | p<0.01 |
|  |  | KO |  |  |  |  |  |  |  |  |  |  | NS | NS |
|  | Runx1 HET | WT |  |  |  |  |  |  |  |  |  |  |  | p<0.01 |

| Inner wall |  | Six1 | Smad4 WT |  |  |  |  |  |  | Smad4 HET |  |  |  |  |
| --- | --- | --- | --- | --- | --- | --- | --- | --- | --- | --- | --- | --- | --- | --- |
|  |  |  | Runx1 WT |  |  | Runx1 HET |  |  | Runx1 KO | Runx1 WT |  |  | Runx1 HET |  |
|  |  |  | WT | HET | KO | WT | HET | KO | WT | WT | HET | KO | WT | HET |
| Smad4 WT | Runx1 WT | WT |  | NS | NS | NS | p<0.01 | p<0.01 | p<0.01 | NS | p<0.01 | p<0.01 | NS | p<0.01 |
|  |  | HET |  |  | NS | NS | NS | p<0.01 | p<0.01 | NS | NS | NS | NS | p<0.01 |
|  |  | KO |  |  |  | NS | NS | p<0.05 | p<0.01 | NS | NS | NS | NS | p<0.05 |
|  | Runx1 HET | WT |  |  |  |  | NS | p<0.01 | p<0.01 | NS | NS | p<0.05 | NS | p<0.01 |
|  |  | HET |  |  |  |  |  | NS | p<0.01 | NS | NS | NS | NS | NS |
|  |  | KO |  |  |  |  |  |  | p<0.01 | p<0.05 | NS | NS | p<0.05 | NS |
|  | Runx1 KO | WT |  |  |  |  |  |  |  | p<0.01 | p<0.01 | p<0.01 | p<0.01 | p<0.01 |
| Smad4 HET | Runx1 WT | WT |  |  |  |  |  |  |  |  | NS | NS | NS | p<0.01 |
|  |  | HET |  |  |  |  |  |  |  |  |  | NS | NS | NS |
|  |  | KO |  |  |  |  |  |  |  |  |  |  | NS | NS |
|  | Runx1 HET | WT |  |  |  |  |  |  |  |  |  |  |  | p<0.01 |

NS: not significant

### Regulatory elements of ΔNp63

The gene-dose-dependent effect of *Six1, Runx1* and *Smad4* on ΔNp63 activation suggests collaboration between these transcription factors in the vaginal cell fate commitment of MDE. The analysis of evolutionally conserved regions by ECR browser [22] identified numerous numbers of putative enhancer elements within *TP63/Trp63* locus. Many of these conserved sequences near ΔNp63 TSS contained binding sites for SMADs, RUNX1 as well as SIX1 (S1 Fig). The 5’ sequence proximal to ΔNp63 TSS, to which SMAD4 binds in VgE but not UtE, also contained binding sites of SMAD4, RUNX1 and SIX1 (S2A Fig). Thus, we genetically tested if the putative 5’ proximal enhancer and the promoter (mm10 Chr16: 25801055-25802045) are sufficient to replicate the expression patterns of ΔNp63 by generating transgenic mice (S2 Fig). However, the transgene (Cre-ires-EGFP) was not expressed in any tissues of 5 founders of transgenic mice. Furthermore, the progenies of the founders carrying *ROSA^mT-mE^* [23] and ΔNp63-Cre-ires-EGFP alleles were also totally negative for EGFP/mEGFP (not shown), indicating the insufficiency of the sequence by itself to replicate the expression patterns of ΔNp63 in MDE. Surprisingly, ΔNp63-Cre knock-in (KI) mice, in which the coding sequence in the first exon of ΔNp63 was replaced with Cre [24] also failed to express the Cre transgene in VgE: When ΔNp63-Cre KI mice were crossed with *ROSA^mT-mE^* reporter mice, the most epithelial cells in vagina of *ROSA^mT-mE^;*ΔNp63*-Cre* KI double-transgenic mice were negative for mEGFP (n=3, S2C Fig). ConTra v3 analysis [25] identified conserved binding sites of SMAD1, SMAD4 and RUNX1 in the sequence deleted in the genome of ΔNp63-Cre KI mice (S1E Fig). Thus, the efficient activation of ΔNp63 locus in MDE appeared to require cooperation of multiple regulatory elements including the protein coding sequence within exon 1. Notably, conserved binding sites of SIX1, RUNX1 and SMADs were not always clustered together, suggesting that these transcription factors may independently act on different enhancer elements. Thus, the gene-dose effects of *Six1, Runx1* and *Smad4* on ΔNp63 activation may reflect the number of active enhancer-like elements that cooperate in the remodeling of ΔNp63 locus.

### Diethylstilbestrol (DES) inhibits activation of ΔNp63 locus in MDE through down-regulation of RUNX1 and SIX1

Previously, we demonstrated that down-regulation of RUNX1 is involved in the pathogenesis of DES-associated vaginal adenosis. DES down-regulated RUNX1 in MDE of vaginal fornix within 24 hours (Fig 6A). However, the effect of 24-hour DES-treatment was more prominent on SIX1 than RUNX1: Nuclear expression of SIX1 disappeared from the MDE in the vaginal fornix and the cervical canal of DES-treated mice (Fig 6A and 6B). DES slightly reduced pSMAD1/5/9 in the vaginal mesenchyme, while DES had no evident effect on the epithelial pSMAD1/5/9 (Fig 6C). In contrast, DES-treatment consistently increased MAPK1/3 activity in vaginal epithelium and mesenchyme (Fig 6C). Hence, the down-regulation of SIX1 was not likely due to the repression of BMP4 or FGF7/10 activity in MDE by DES.

**Fig 6.**
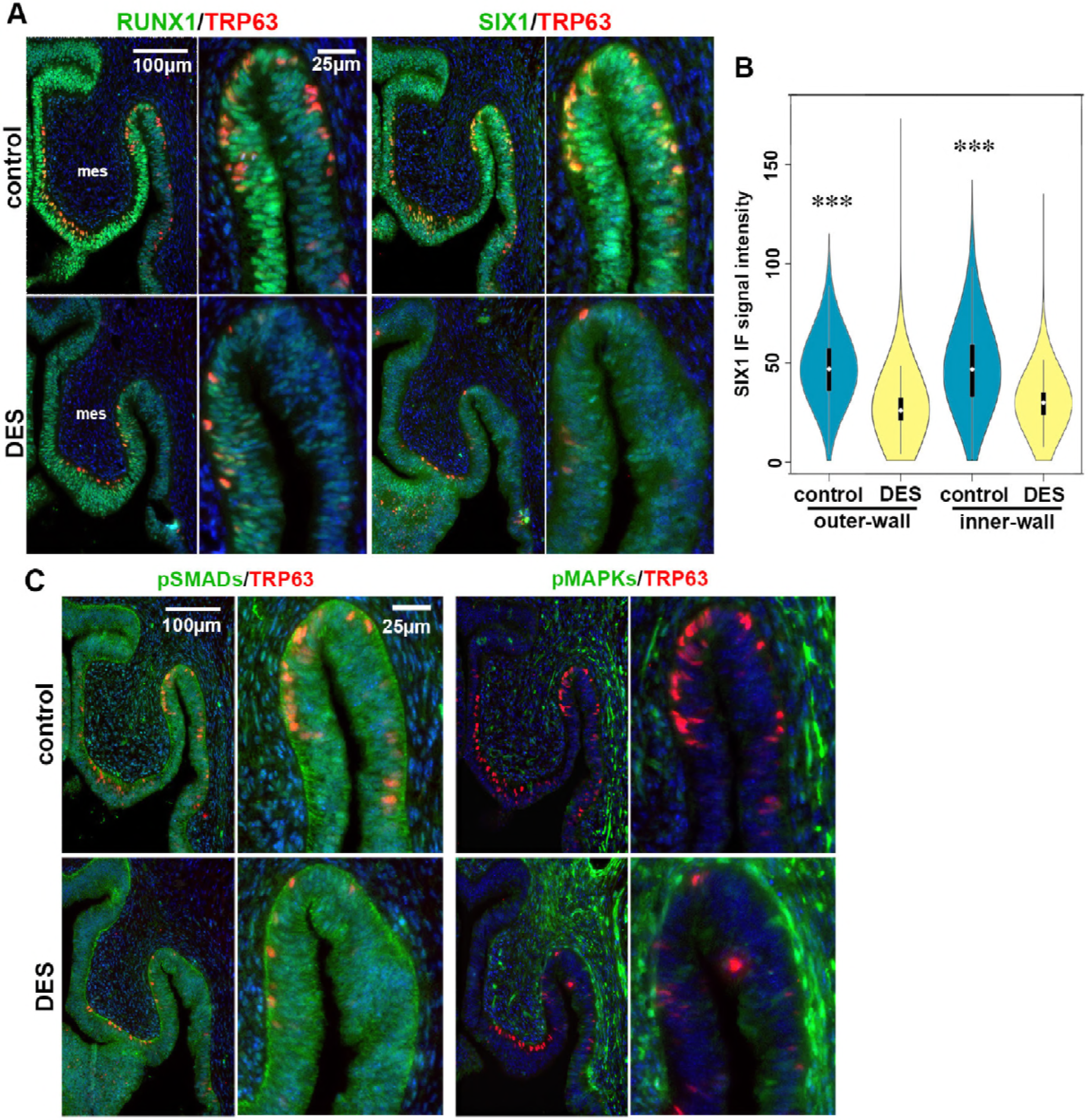
DES inhibits expression of SIX1 and RUNX1 in the vaginal fornix. (A-C) IF analysis for DES-effects on essential factors in the activation of ΔNp63 locus in MDE. mes; mesenchyme. FRTs are collected from PD2 female mice with/without DES treatment (24 hours after DES-pellet injection). (A) IF assay of SIX1 and RUNX1. (B) Violin plot presentation of SIX1 IF signals in the outer and inner fornix walls of control and DES-treated PD2 mice (n=4 each). SIX1 IF signals in MDE were significantly higher (*** p< 0.01) in control than DES-treated mice in both outer and inner fornix walls. (C) IF assay of pSMAD1/5/9 and pMAPK1/3.

Our previous tissue recombination study has established that DES blocks expression of ΔNp63 in MDE through estrogen receptor *α* (ESR1) within epithelial cells [4, 8]. The expression patterns of SIX1, RUNX1 and ΔNp63 in the fornix of *Esr1* cKO mice were indistinguishable from these in wild type mice at PD3 (Fig 7A). In agreement with the tissue recombination study, DES did not block the induction of ΔNp63 in the VgE when *Esr1* was deleted in MDE by *Wnt7a-Cre* (*Esr1* cKO mice) (Fig 7B). Moreover, DES exposure promoted the expression of ΔNp63 in VgE in *Esr1* cKO mice (Fig 7B), forming a continuous layer of ΔNp63-positive cells by PD3, ≥ 1 day earlier than normal development. Therefore, ESR1 in epithelium and mesenchyme has the opposite effect on the induction of ΔNp63 (S3 Fig). In fact, DES-treatment induced RUNX1 and SIX1 in the UtE of *Esr1* cKO mice by PD3 (not shown). DES-ESR1 activity attenuates the expression of SIX1 and RUNX1 in MDE cell-autonomously, as the expression of SIX1 and RUNX1 was maintained in the vaginal fornices of *Esr1* cKO mice (Fig 7C). Interestingly, the effect of DES on MAPK1/3 activity in MDE and mesenchyme was exaggerated in the vagina of *Esr1* cKO mice (Fig 7C), indicating that epithelial ESR1 repressed the FGF7/10-MAPK1/3 signaling pathway in both vaginal epithelium and mesenchyme. A continuous layer of ΔNp63-positive basal cells formed by PD3 on the outer wall of vaginal fornix in MAP2K1^DD^ conditional transgenic mice (Fig 7D, control), suggesting that DES promotes vaginal differentiation of MDE by activating the MAPK pathway. Nevertheless, DES repressed the expression of SIX1 and ΔNp63 in MDE expressing MAP2K1^DD^ (Fig 7D, DES). Thus, the activation of MAPK pathway alone did not protect MDE from DES-induced vaginal adenosis.

**Fig 7.**
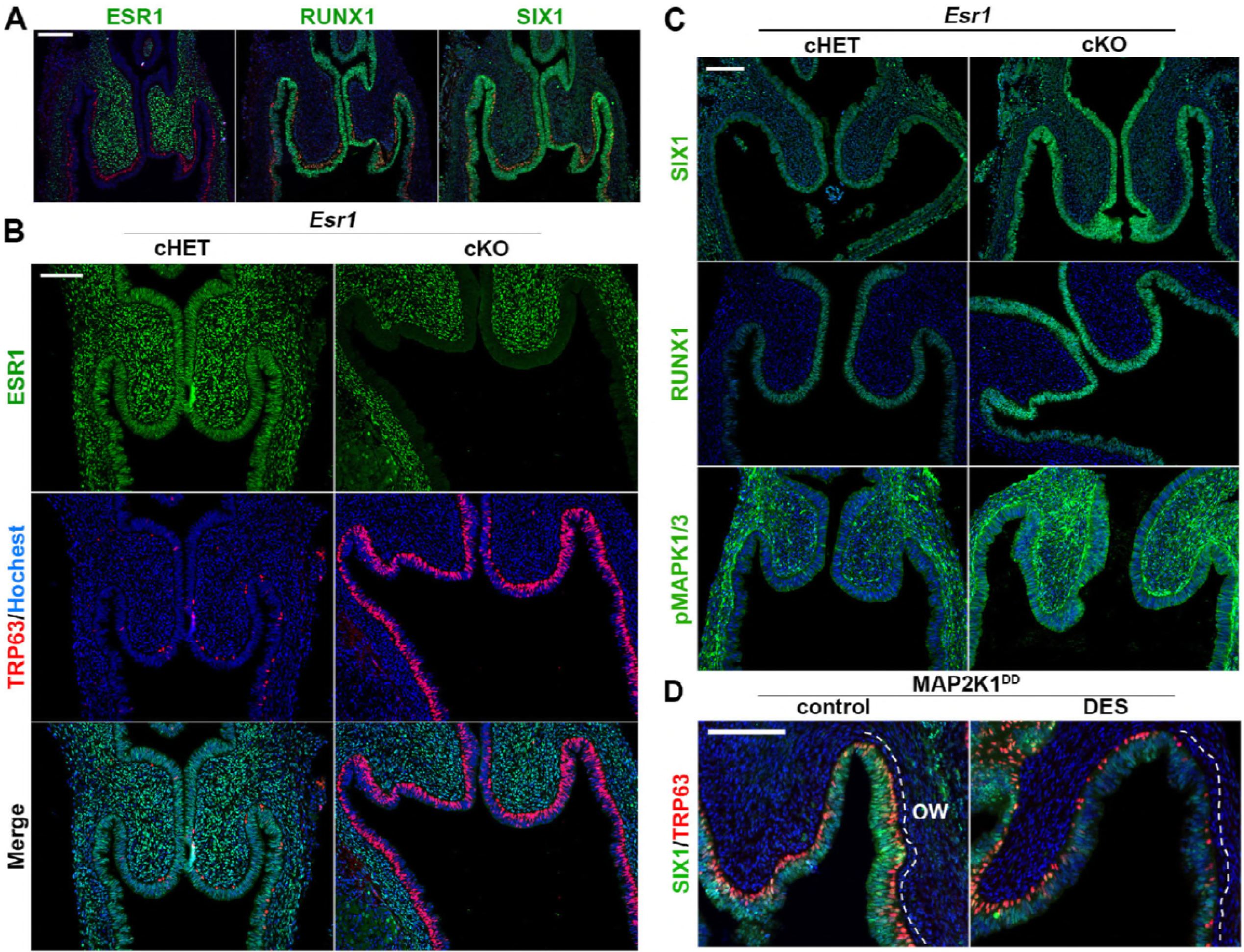
Epithelial ESR1 mediates DES effects on ΔNp63 in developing vagina. (A) Expression patterns of ESR1, RUNX1 and SIX1 in *Esr1* cKO mice (PD3) are indistinguishable from wild type mice. (B and C) Effect of DES on the FRT of *Esr1* cHET and cKO mice (PD3): (B) IF assay of ESR1 (green) and TRP63 (red), (C) IF assay of RUNX1 and pMAPK1/3. (D) Effect of DES on the expression of SIX1 (green) and TRP63 (red) in the fornix of *Map2k1^DD^* conditional transgenic mice (PD3). The outer-wall (ow) of fornix is marked with dotted lines. Bars = 100 μm.

## DISCUSSION

It has long been known that the differentiation of MDE into distinctive epithelia of uterus and vagina is under the control of organ-specific mesenchyme [5]. Through a series of studies with genetically engineered mice, our group has established that ΔNp63 is the master regulator of vaginal epithelial differentiation in MDE [8], and that the expression of ΔNp63 is induced by mesenchymal paracrine factors, BMP4, ActA and FGF7/10 [11, 12]. Within MDE, the signals from underlying mesenchyme are transduced by BMP4-SMADs, ActA-RUNX1 and FGFs-MAPKs. Since mouse vaginal mesenchyme can induce ΔNp63 and squamous differentiation in human MDE, the molecules that mediate the communication between mesenchyme and epithelium in the commitment of MDE to vaginal cell fate must be common between these two species [26].

In this study, we identified SIX1 as a key transcription factor that mediates the mesenchymal signals in the activation of ΔNp63 locus during vaginal cell fate commitment of MDE. Subsequently, we propose that vaginal mesenchymal factors induce MDE to commit to vaginal epithelial cell fate by activating ΔNp63 locus through cooperation of multiple enhancer elements, which are activated by SMADs, RUNX1 and/or SIX1 (Fig. 8). An enhancer is a genomic region of few hundred base pairs that contains clustered binding-sites for multiple transcription factors. Although many transcription factors cannot bind their target site in the context of nucleosomal DNA, enhancer-mediated simultaneous-binding of multiple transcription factors can overcome the nucleosome barrier [27]. Thus, enhancers integrate multiple signaling pathways through binding of downstream effectors [28, 29]. In cell fate commitment of MDE to VgE, BMP, ActA and FGF pathways are integrated to prime VgE-specific gene expression programs in MDE through the simultaneous binding of SMADs, RUNX1 and SIX1 to ΔNp63 enhancers (Fig. 8). Approximately 80% of all characterized mouse enhancers show tissue-specific expression [30]. In this regard, the enhancers that regulate ΔNp63 expression in MDE must be distinctive from those in the skin because *Six1* null [19] and *Runx1* null [31] mice do not exhibit the deformation of skin and appendages observed in ΔNp63 mutant mice [32]. The identification of key regulator elements of ΔNp63 in MDE is imperative to fully appreciate the pathogenesis of vaginal adenosis, which is a result of faulty cell fate commitment of VgE. However, enhancers can regulate the expression of genes that are mega-bases apart [30, 33]. Therefore, the identification of key regulator elements of ΔNp63 in MDE requires genome-wide screening of transcription factor binding sites by chromatin immunoprecipitation-sequencing (ChIP-seq). However, the usage of ΔNp63 enhancers must be unique between different regions of MDE as demonstrated by the difference in the requirement of SMAD4, RUNX1 and SIX1 for ΔNp63 expression in mouse genetic studies. Given the heterogeneity of the cell population, the narrow developmental time window, and the small tissue amount of MDE, the identification of ΔNp63 regulatory elements in MDE by current standard techniques is challenging.

**Fig 8.**
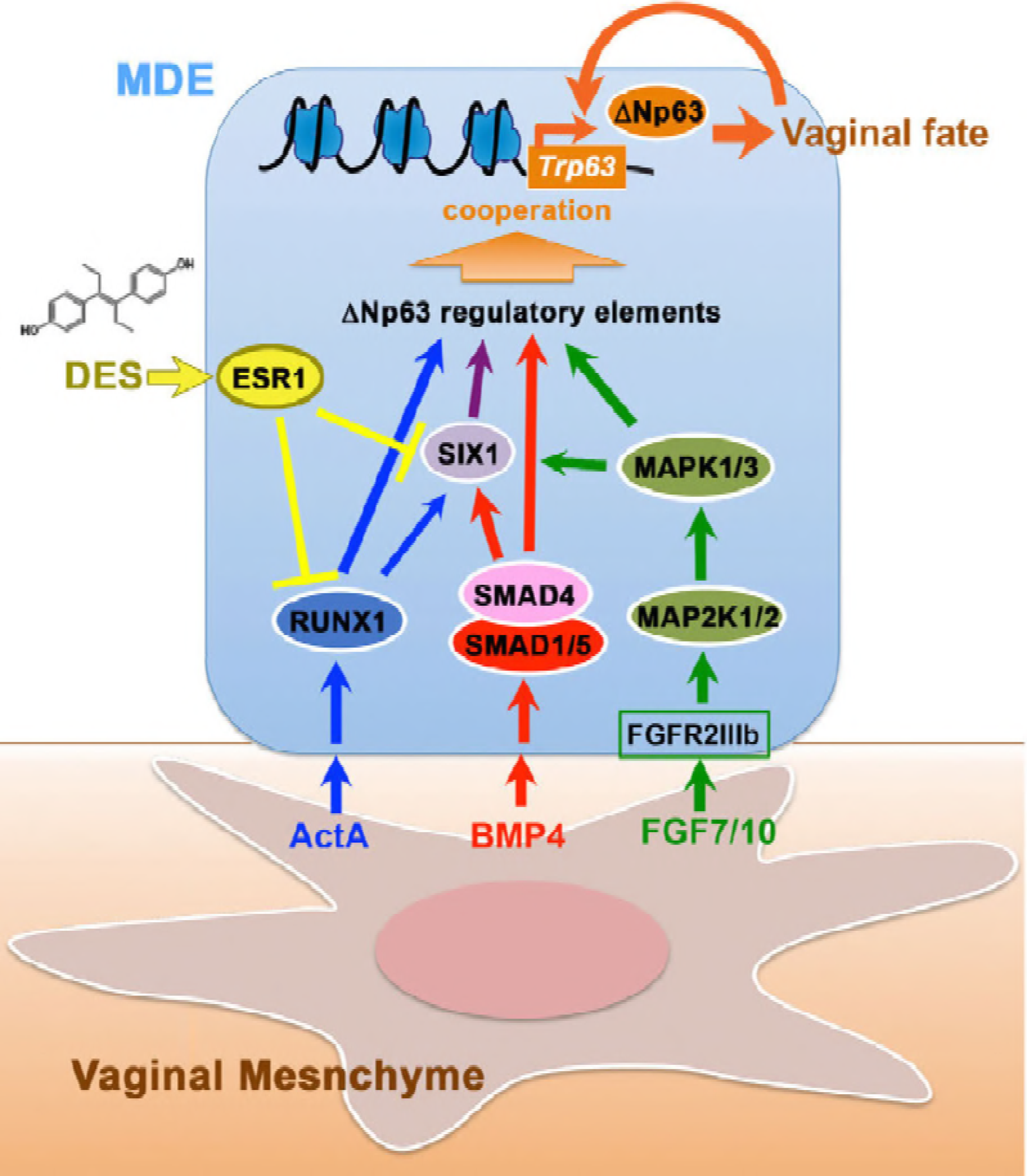
Model of vaginal epithelial cell fate commitment in MDE. Signals of vaginal mesenchymal factors are transduced to down-stream transcription factors, and the transcription factors dose-dependently activate enhancers of ΔNp63 in MDE. Upon differentiation of VgE, ΔNp63 itself maintains the transcriptional activity of ΔNp63 locus in VgE fate in dependent of vaginal mesenchymal factors. DES-ESR1 activity within MDE causes vaginal adenosis by blocking the vaginal cell fate commitment of MDE interfering the signal transduction. Meanwhile, DES-ESR1 activity in vaginal mesenchymal cells promote activation of ΔNp63 locus in MDE through paracrine mechanisms.

Most vaginal adenocarcinomas (VACs) are believed to arise from vaginal adenosis because of the presence of adenosis lesions at the primary site of VACs. Hence, better understanding in etiology of vaginal adenosis is particularly crucial in order to develop preventive and therapeutic approaches for VACs. In the past, *in utero* DES-exposure was the primary cause of vaginal adenosis and VAC. Since the expression patterns of ΔNp63 and RUNX1 as well as the effect of DES on the expression of these transcription factors are identical between human and mouse MDE [1, 10, 26, 34], the molecular model established in mice (Fig 8) should explain the etiology of vaginal adenosis in DES-exposed women. However, VACs occur in women who have no history of DES exposure [16, 35]. Because the expression of ΔNp63 in the lower MDE occurs during the first trimester in human fetus [10, 34], the pathogenesis of non-DES-associated VACs should still involve an *in utero* event that disturbs the vaginal epithelial cell fate commitment in MDE. In this regard, exposure to a compound that inhibits any pathways/molecules described in Fig 8 can lead to vaginal adenosis. As cell fate commitment of mouse VgE occurs in the first week of postnatal development [3, 10], neonatal mice would be useful to screen medical and environmental chemicals that interfere ΔNp63 expression in VgE.

Some studies suggest the *de novo* formation of adenosis in the vagina of adult women associated with medical treatments [36-38]. However, given the low detection sensitivity of routine colposcopy and cytology screenings for adenosis, adenosis cases that appear to be *de novo* are likely due to an increased visibility of previously imperceptible adenosis lesions enlarged by a reactive change to medical treatments.

In addition to vaginal adenosis, perinatal DES exposure of female mice induces uterine squamous metaplasia [39], a formation of squamous epithelium within the simple columnar UtE. Interestingly, the gene expression pattern of uterine squamous metaplasia lesions is identical to that of normal VgE [4, 8], indicating that uterine squamous metaplasia is vaginal cell fate commitment of MDE within the uterus. Thus, developmental DES exposure elicits opposite effects on the cell fate commitment of MDE in developing uterus versus vagina. This intriguing dual-effect of DES is explained by the opposite functions of epithelial versus mesenchymal ESR1. As shown in our current study, DES action through epithelial ESR1 interferes the activation of ΔNp63 locus whereas DES action through mesenchymal ESR1 promotes ΔNp63 expression (S3 Fig). When ESR1 is expressed in both epithelium and mesenchyme, DES effects via epithelial ESR1 are dominant. In developing uterus and vagina, ESR1 is initially expressed only in mesenchymal cells, and ESR1 expression in MDE occurs at the posterior end and gradually progresses anterior [7, 34]. DES exposure most efficiently induces vaginal adenosis at the first trimester of human fetuses and the postnatal day 1-5 in neonatal mice, when ESR1 is expressed in the MDE of the vagina but not the uterus. Accordingly, DES blocks ΔNp63 activation in the vagina through epithelial ESR1 and activates ΔNp63 in the uterus through mesenchymal ESR1. Our current study predicts that DES induces expression of BMPs, FGFs and Activin/TGFβ through ESR1 in the uterine mesenchyme. On the other hand, molecular mechanisms through which DES represses RUNX1 and SIX1 in MDE remain unclear. Elucidating the underlying molecular pathogenesis of DES-associated adenosis will help identify etiology of non-DES-associated vaginal adenosis and VAC.

## METHODS

### Mouse models

All animal procedures were approved by the Animal Care and Use Committee in the Ohio State University. The mouse strains carrying the following alleles were utilized: *Six1^flox^* [*Six1^tm2^*^.1^*^Mair^*] [40], *Trp63^flox^* [*Trp63^tm3.2Brd^*] [41], ΔNp63-EGFP knock-in (*Trp63* ^ΔNp63^*^-EGFP-KI^*) [32], *Runx1^flox^* [*Runx1^tm1Tani^*] [42], *Fgfr2^flox^* [*Fgfr2^tm1Dor^/J*] [43], *ROSA^mT-mE^* [*Gt*(*ROSA*)*26Sor^tm4^*^(^*^ACTB-tdTomato,–EGFP^*^)^*^Luo^/J*] [23], *ROSA^MAP2K1DD^* [*Gt*(*ROSA*)*26Sor^tm8^*^(^*^Map2k1*,EGFP^*^)^*^Rsky^/J*] [44], *Smad4^flox^* (*Smad4^tm2.1Cxd^/J*) [45], *Esr1^flox^* [46], *Bmp4^flox^* [*Bmp4^tm1Jfm^*] [47], *Twist2^Cre^* [*Twist2^tm1.1^*^(^*^cre^*^)^*^Dor^*] [17], *Pax2-Cre* [Tg(Pax2-cre)1Akg] (MMRRC) [48], *Wnt7a-Cre* [20] and ΔNp63-Cre [*Trp63^tm1.1^*^(^*^cre^*^)^*^Ssig^/J*] [24]. C57BL/6J mice were purchased from Jackson Laboratory (Bar Harbor, ME). MDE-specific conditional knockout (cKO) and conditional heterozygous (cHET) mice were generated by crossing lines carrying floxed alleles with *Wnt7a-Cre* mice, except for *Trp63^flox^* mice, which were crossed with *Pax2-Cre*. *Twist2^Cre^* mice were used for mesenchyme-specific deletion of *Bmp4*. The day of birth was count as PD1.

### Neonatal DES treatment

A ~40 μg DES slow-release pellet was prepared as previously described [12]. The ~0.04 mg/mm DES filled tubing was cut into 1 mm length and subcutaneously injected into newborn mice using a 19 gauge trocar.

### Immunofluorescence (IF) and immunohistochemistry (IHC)

IF and IHC assays were performed as previously described [49]. Briefly, tissues were fixed with Modified Davidson’s fixative solution (Electron Microscopy Sciences, Hatfield, PA), processed into paraffin, and sectioned at 5 μm. The sections were heated in 10 mM sodium citrate buffer (pH 6.0) containing 0.05% Tween-20 for 35 min in an Electric Pressure Cooker. The following primary antibodies were used at the indicated dilutions: anti-CTNNB1 (CAT-5H10) (1:100, 13-8400) from ThermoFisher (Waltham, MA); anti-TRP63 (4A4) (1:200, 790-4509) from Ventana Medical Systems (Tucson, AZ); anti-ΔNp63 (1:2,000, PC373) from Millipore (Billerica, MA); anti-PGR (1:200, A0098) from Agilent Technologies (Santa Clara, CA); anti-RUNX1 (2593-1, 1:400) from Epitomics (Burlingame, CA); anti-phospho (p)-MAPK1/3 (p-T202/Y201, 1:30, #4370) and anti-pSMAD1/5/9 (1:50, #9511) from Cell Signaling Technology (Danvers, MA); anti-GFP (1:100, ab6673) from Abcam (Cambridge, MA); anti-SIX1 (1:800, HPA001893) from Sigma-Aldrich (St. Louis, MO); anti-ESR1 (1:100, RM-9101) from Lab Vision (Fremont, CA). For IF assay, Alexa-Fluor594 anti-mouse IgG (H+L) (1:1,000, 715-586-150) and Alexa-Fluor488 anti-rabbit IgG (H+L) (1:1,000, 711-546-152) from Jackson ImmunoResearch (West Grove, PA) were used for the secondary antibodies, and bisbenzimide H 33258 (Hoechst 33258, 1:10,000, Sigma-Aldrich) was used for nuclear staining. For IHC with DAB (3,3’-diaminobenzidine, Sigma-Aldrich), biotinylated anti-rat IgG (H+L) (1:800, 712-066-153) was used in conjunction with streptavidin-horseradish peroxidase (1:400, 016-030-084, Jackson ImmunoResearch). Micrographs were captured using a BZ-9000 microscope (Keyence, Osaka, Japan) under identical conditions between samples for each antibody. The contrast of images was adjusted by applying identical parameters to the images for each antibody with the batch-process function of Adobe Photoshop CS6 (Adobe, CA, San Jose, CA, USA). To capture a wide area in a single image, tissue sections were scanned in multiple frames, and the images were automatically merged together utilizing the Image Stitching function of image analysis tool.

### Morphometric analysis

The methods for the quantitative analysis on the squamous transformation of MDE [12] and the IF signal [50] were previously described. We adapted these methods with some modifications. The length of epithelium at the basal lamina was measured in the outer-wall of vaginal fornix in at least 2 sections per animal in TRP63 immunostained sections. The proportion of epithelium with ΔNp63-psotive basal layer was calculated by “length of epithelial basement membrane associated with TRP63-positive cells” ÷ “total epithelial basement membrane length” × 100, for each mouse.

Basal cell density in the outer and inner fornix walls was calculated by number of TRP63-positive pixels per epithelial basement membrane length. In tissue sections of vaginal fornices stained for TRP63, epithelial areas were manually selected, and the pixels positive for TRP63 signal within the epithelium were selected by adjusting the lower threshold for positivity to exclude background noise. Epithelial basement membrane was manually marked on the IF images, and the p63-positive area and the basement membrane length were measured utilizing Image J (NIH, Bethesda, MD). Analysis was performed on ≥ 4 fornice from ≥ 3 mice per group. The value in each fornix was considered as a single measurement. Statistical significance was analyzed by One-way ANOVA with post-hoc Tukey’s HSD Test.

### SIX1 IF analysis

Quantitative IF assay was performed as previously described with modifications [50]. Tissue sections for an analysis were stained together, and images were captured at the same time under the identical conditions. Images of ≥ 4 tissue sections from n≥ 3 independent animals were analyzed for each group. Epithelial areas were manually selected, and the signal intensity per pixel within the epithelial area was measured by Image J. In all experiments, approximately equivalent areas were analyzed in each sample, and there was no significant intragroup difference in the average signal intensity. Thus, all samples in each group were plotted together, and the distributions of signals were compared between groups by the Mann–Whitney ⋃ test with continuity correction.

### Immunoblot analysis

Ovaries, uteri and vaginae from PD2 mice (5-6 mice per blot) were homogenized with a minipestle in ice-cold lysis buffer containing protease (cOmplete Protease Inhibitor Cocktail,Roche) and phosphatase (phoSTOP, Roche) inhibitors and loaded onto NuPAGE 4–12% Bis-Tris precast SDS-PAGE gel. Proteins were transferred to a PVDF membrane (Millipore Sigma, St. Louis, MO, USA). The membrane was incubated with anti-RUNX1 antibody (1: 2000, Epitomics), anti-SIX1 antibody (1:1,000, Millipore Sigma) and GAPDH (1: 2000, G8795, Millipore Sigma) in the OdysseyR Blocking buffer (TBS) (from LI-COR Biosciences, NE, USA) overnight at 4 °C. IRDye^®^ 800CW Donkey anti-rabbit IgG, IRDye^®^ 680LT Donkey anti-rabbit IgG and IRDye^®^ 680LT Donkey anti-mouse IgG were used for the secondary antibodies. The signal was detected using Odyssey CLx Imaging System (LI-COR Biosciences, NE, USA). The analysis was repeated 3 times with independent samples.

### Uterine organ culture

Uterine hanging drop organ culture was performed as previously described with minor modifications [11]. Briefly, uteri were dissected from PD1 mice, cleaned by removing connective tissues, and cut into 3 pieces per uterine-horn in Dulbecco’s Modified Eagle Medium/Nutrient Mixture F-12 (DMEM/F12, 11039, Life Technologies) containing 10 nM ICI 182,780 (Sigma-Aldrich). The uterine pieces were then placed in autoclaved PCR tube caps (AXYGEN, Union City, CA) with basal medium (10 nM ICI 182,780 DMEM/F12 with Insulin-Transferrin-Selenium and Antibiotic-Antimycotic) with/without 20 ng/ml human recombinant BMP4, ActA and/or FGF10 (Life Technologies), inverted, and incubated. Uterine pieces were cultured up to 3 days with daily medium change, fixed with Modified Davidson’s fixative, and processed for histological analysis.

## ACKNOWLEDGMENTS

The authors thank Dr. Altea Rocci, Shayna Wallace, Justin Thomas and the Solid Tumor Biology Research group histology core for technical help. This work was supported by the National Institutes of Health [RO1CA154358, RO1HD064402, P30CA016058 to T.K.].

## Supporting information captions

**S1 Fig.**
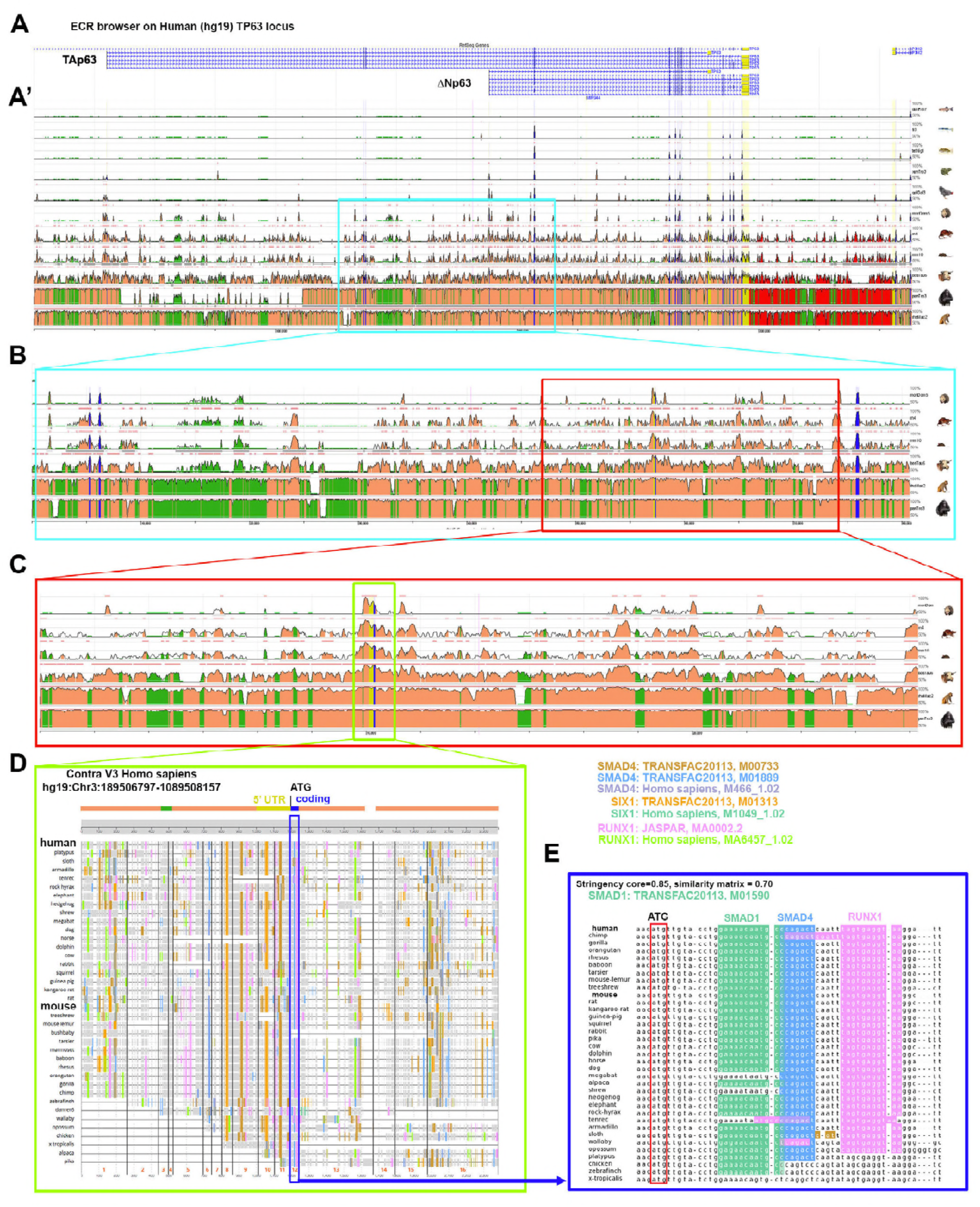
(A-C) ECR browser view of *Homo sapiens TRP63*. (D) Contrav 3 analysis. Conserved SMADs:RUNX1 binding sites in the coding region in ΔNp63 exon 1. A boxed region with a colored line in each panel is enlarged in the next panel. (E) The sequence is deleted in the ΔNp63-Cre KI mouse genome.

**S2 Fig.**
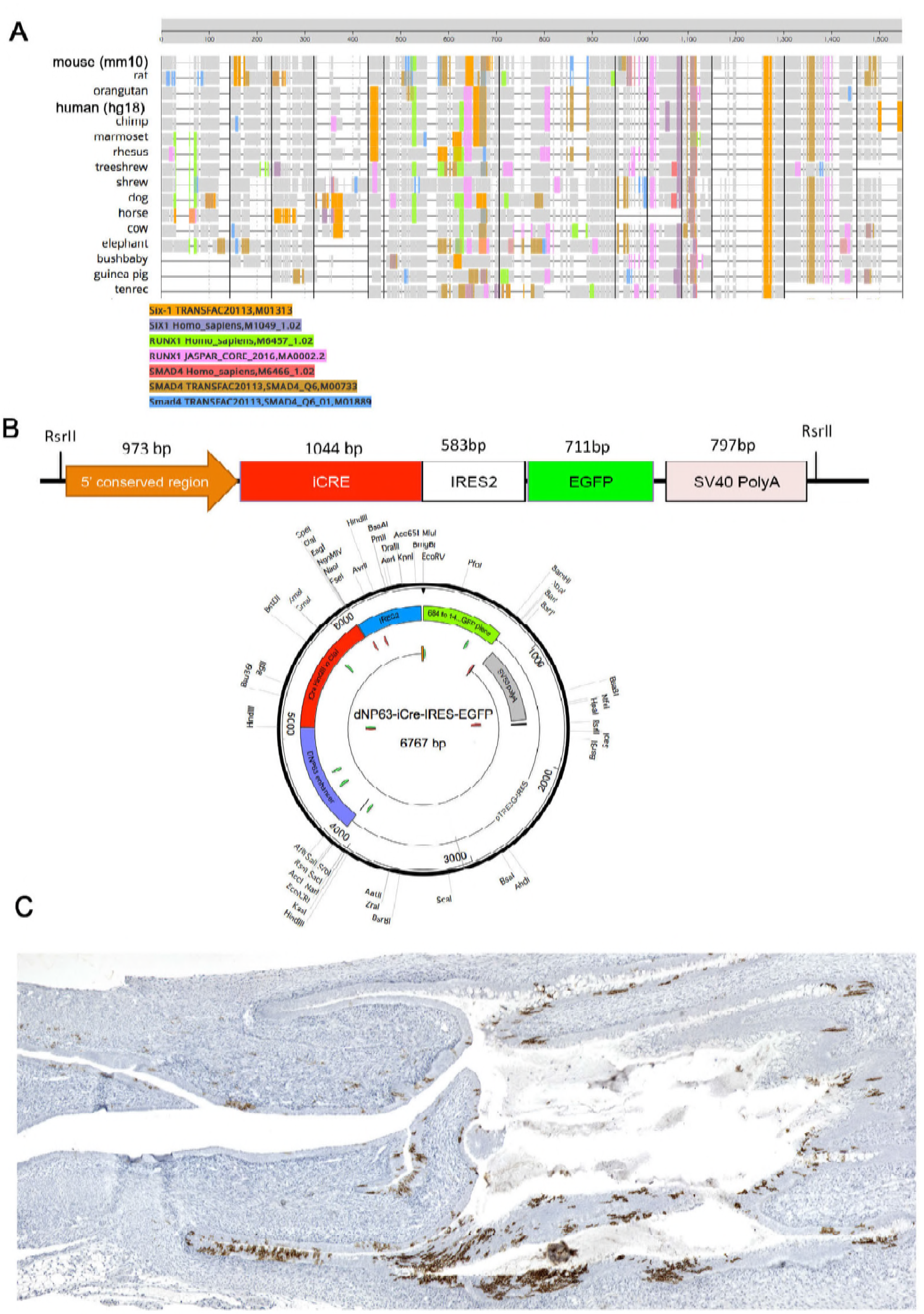
Structure of ΔNp63-iCre-IRES-EGFP transgene. (A) Contra V3 analysis of the putative 5’ proximal enhancer (based sequence: mm10 chr16:25801055-25802045). (B) Vector maps of the ΔNp63-iCre-IRES-EGFP transgene (linearized and circular form). (C) Distribution of cells expressed ΔNp63-Cre in the lower FRT of PD21 mice. mEGFP reporter (brown) is detected by IHC.

**S3 Fig.**
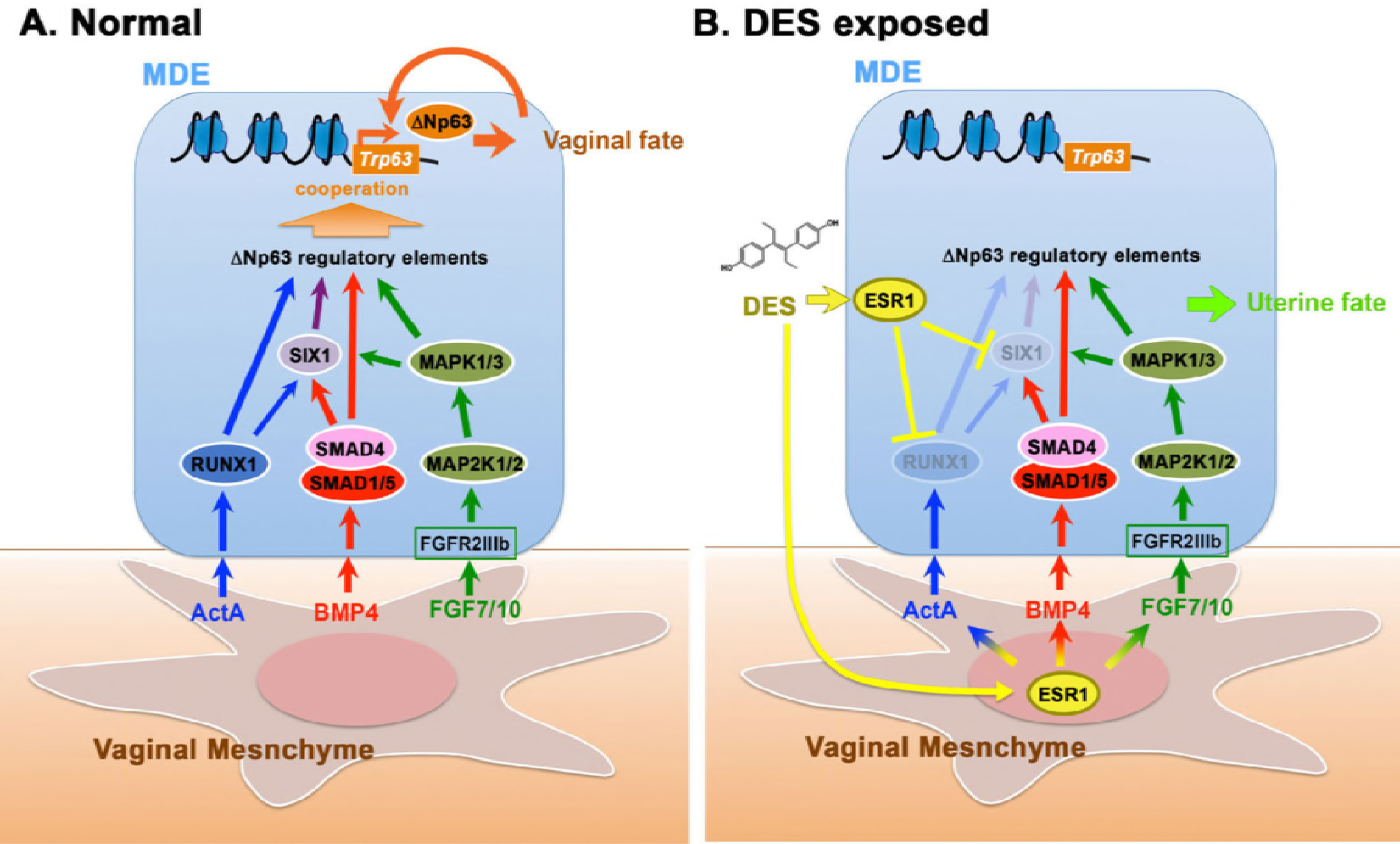
Model: Effect of DES exposure on the signaling pathways in developing vagina.

